# Age-Related Changes in Curiosity: The Influence of Locus Coeruleus on Information-Seeking Behavior

**DOI:** 10.1101/2025.09.04.674248

**Authors:** Hsiang-Yu Chen, Emma Lepore Carlson, McKenna S. Costello, Johanna L. Matulonis, Alex A. Adornato, Katherine E. O’Malley, Jacob M. Hooker, Heidi I. L. Jacobs, Anne S. Berry

## Abstract

Curiosity enhances learning and memory and has been linked to the locus coeruleus (LC), which undergoes age-related decline. To examine how aging affects curiosity and information-seeking, we developed the Photographic Art Storytelling Task. Participants (sixty-eight young and sixty-five older adults) viewed photographs, rated their curiosity, and later read stories associated with selected images. The stories were deliberately constructed to be either interesting or boring, functioning as rewards that elicited prediction errors. Participants reappraised their curiosity, allowing us to separate intrinsic motivation from story-driven reward influences. Associations between performance and both pupil diameter and MRI measures of LC integrity supported a role of LC in curiosity regulation. Aging was associated with greater reliance on novelty-driven (initial) curiosity and a shift away from prediction error-related information-seeking. Both age groups showed curiosity-driven memory enhancement, with older adults exhibiting youth-like effects, suggesting a role for intrinsic motivation in preserving cognitive function in aging.

## Introduction

Curiosity is a fundamental motivation that drives individuals to seek new knowledge. When effectively engaged, it can enhance both learning and memory. But how is curiosity ignited to support information-seeking behavior? Research suggests that two neuromodulatory systems are involved: the dopamine system, which is linked to prediction errors, and the norepinephrine system, which responds to novelty. Gruber and Ranganath (2019)^1^ proposed a comprehensive framework that explains how curiosity enhances memory, highlighting the interaction between brain mechanisms, cognitive processes, behavioral responses, and individual perceptions during learning. According to their model, curiosity arises from context-dependent prediction errors, which refer to discrepancies between new information and existing knowledge or expectations. This gap in understanding motivates individuals to seek additional information and enhances memory consolidation. The resolution of uncertainty is experienced as rewarding, engaging dopaminergic activity in the substantia nigra (SN) and ventral tegmental area (VTA), which in turn influences the hippocampus^2–4^.

In contrast, Takeuchi (2024)^5^ highlighted a complementary role for the LC in curiosity driven by novelty. When new information bears no clear connection to prior knowledge, the LC plays a key role in initiating curiosity. The LC-hippocampal system is implicated in the early phases of memory consolidation in response to novelty. Both animal^6–8^ and computational^9^ studies support this view, showing that LC-norepinephrine neurons become active in exploring novel environments. Together, these findings suggest that curiosity-driven information-seeking can arise from either context-based prediction errors (via the SN/VTA-dopamine system) or exposure to novelty (via the LC-norepinephrine system). Both systems facilitate exploration and learning by enhancing memory consolidation in the hippocampus. Importantly, these cognitive neuromechanisms are not mutually exclusive pathways; rather, they form a continuum, reflecting a dynamic balance between the two neuromodulatory systems in guiding curiosity and learning^5^ (Fig. 1).

**Fig. 1.**
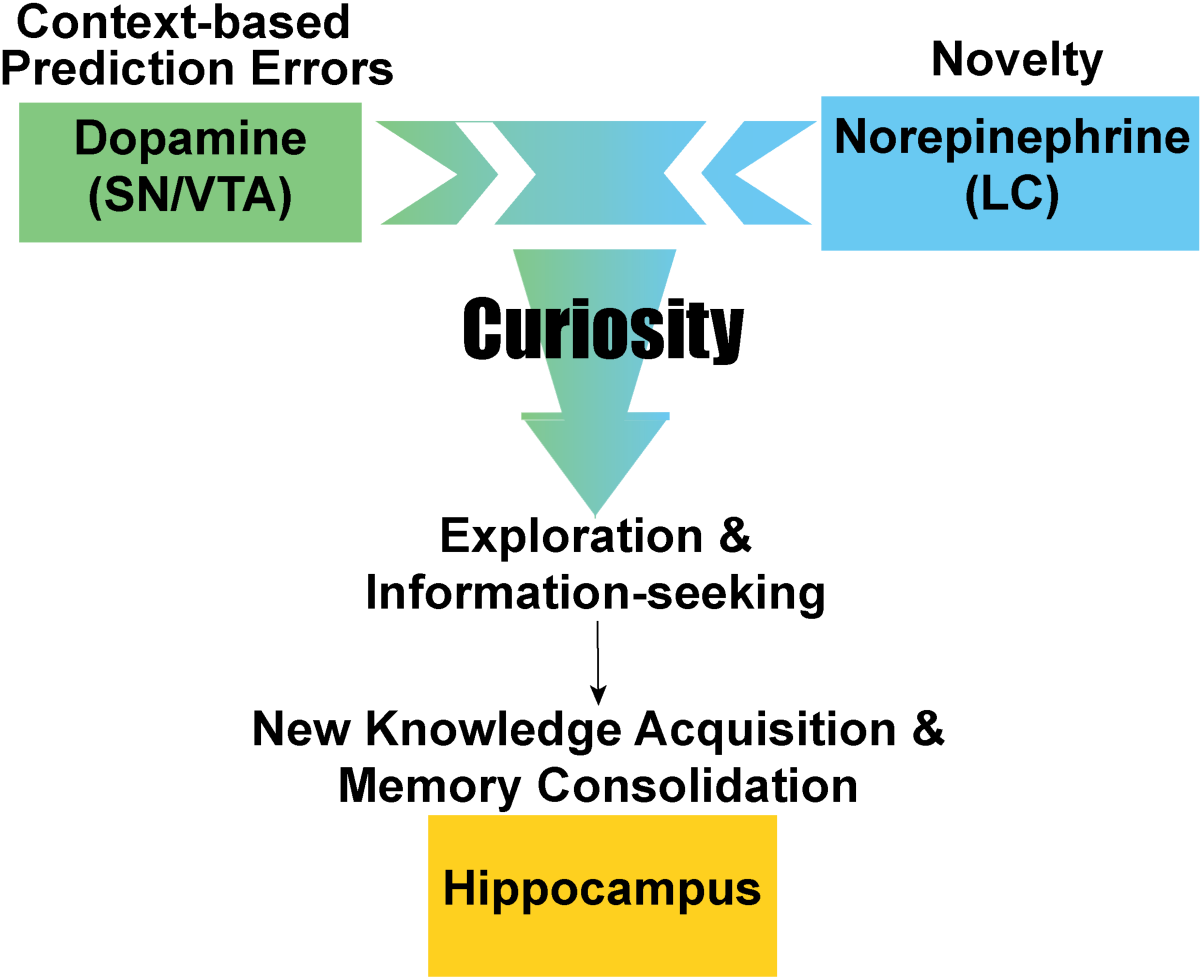
Dual neuromodulatory systems of curiosity-driven information-seeking and memory consolidation. This framework distinguishes between two forms of curiosity: context-based prediction errors, which are associated with the substantia nigra (SN)/ventral tegmental area (VTA) and the dopamine system, and novelty, which is mediated by the locus coeruleus (LC) and the norepinephrine system. Context-based prediction errors refer to the discrepancy between the information acquired and the knowledge already learned. In contrast, novelty occurs when the information acquired is unrelated to prior knowledge. Both forms elicit curiosity to explore and seek information. The acquired knowledge is consolidated as a new memory in the hippocampus. This dual system of curiosity-driven information-seeking exists along a continuum.

In the course of healthy aging, both dopamine and norepinephrine catecholamine systems undergo degeneration, which affects various cognitive functions. Specifically, when motivated learning involves extrinsic rewards such as monetary incentives, older adults, compared to young adults, show reduced blood-oxygen-level-dependent (BOLD) signal changes associated with reward prediction errors in the medial prefrontal cortex (mPFC) and nucleus accumbens (NAc)^10,11^. This age-related decline in processing prediction errors is thought to stem from changes in the dopamine system^12,13^. For instance, in a combined fMRI and pharmacological study, Chowdhury and colleagues (2013)^14^ found that older adults who received a dopamine-enhancing drug demonstrated improved learning, along with activation in the NAc during a monetary reward decision-making task, activation typically associated with reward prediction errors. These findings suggest an age-related decline in prediction error processing, which might influence curiosity-driven information-seeking.

While much attention has focused on dopaminergic decline, emerging evidence points to the LC, the brain’s primary source of norepinephrine, as another key player in cognitive aging. Recent advances have enabled in vivo assessment of LC structure using magnetic resonance imaging (MRI), revealing a quadratic trajectory of LC-MRI contrast across the lifespan, with signal increasing to a peak around age 60, followed by a gradual decline in late adulthood^15–17^. Age-related changes in LC structural integrity have been associated with cortical thickness and memory performance in older adults^18–21^. Individuals with higher LC integrity exhibited better memory performance. However, it remains unclear to what extent aging and LC function specifically influence curiosity-driven information-seeking, and whether such curiosity can enhance learning and memory in late life. Notably, preserved LC function has been associated with cognitive resilience against Alzheimer’s disease-related pathology^15,22–25^, suggesting that the LC-norepinephrine system may help compensate for age-related neural decline. This evidence raises the possibility that LC-driven information-seeking, such as novelty-driven curiosity, could serve as a pathway for supporting learning and memory in older adults. Building on this idea, the present study aims to investigate whether engaging the LC-norepinephrine system through curiosity can enhance learning and memory in late life.

To address how aging influences curiosity-driven information seeking and memory, we developed a novel paradigm—the Photographic Art and Storytelling Task (PAST). This task integrates curiosity and reward contexts with memory assessment to examine both novelty-driven and prediction error-based information-seeking behavior, as well as subsequent memory consolidation. PAST is uniquely suited to generate rich behavioral data, including self-reported curiosity ratings, and allows us to examine how aging may shift the balance between novelty-seeking and prediction error-based exploration. In addition to behavioral measures, we recorded pupillary responses during the task as a psychophysiological marker of curiosity-related arousal^26,27^, and structural MRI was used to assess LC integrity. These multimodal measures enabled us to investigate how curiosity is supported by the LC-norepinephrine system in aging. While both the dopamine system, which supports prediction error-driven learning, and the norepinephrine system, which underlies LC-mediated novelty processing, show age-related changes, emerging evidence suggests that dopamine-related frontostriatal mechanisms decline more steeply with age^10,11^. In contrast, relatively preserved LC-NE integrity in some older adults may represent a mechanism of cognitive resilience^15,22–24^, potentially allowing them to engage more readily with novelty-driven motivation and learning. Given these differential trajectories, we hypothesized that young adults, with relatively intact prediction error processing, would engage more in prediction error-based information-seeking, whereas older adults would rely more heavily on novelty-driven curiosity. Furthermore, individuals with greater LC structural integrity were expected to exhibit more novelty-driven information-seeking. We also predicted that pupil dilation would increase with curiosity levels, reflecting elevated arousal and attentional engagement. This effect was anticipated across age groups, though potentially attenuated in older adults, in line with age-related changes in arousal systems associated with the LC-norepinephrine system. Finally, we examine whether individual differences in curiosity predict behavioral markers of memory performance, with the aim of understanding how motivational factors may contribute to cognitive resilience in aging.

## Results

Sixty-eight young adults and 65 older adults participated in this study. Participants completed the PAST, which comprised several sequential phases (Fig. 2; see the Methods section for full details). Briefly, in Phase 1, participants were shown black-and-white photographs and rated their curiosity about learning the story behind each image. Because participants were unaware of the photographs and their associated stories prior to the task, the curiosity ratings provided in this phase likely reflect novelty-driven curiosity. Based on these ratings, we created 12 high-curiosity and 12 low-curiosity photograph pairs to establish curiosity contexts. In Phase 2, participants were presented with the photograph pairs and asked to choose one image from each pair to reveal the associated story. In Phase 3, participants read part of each chosen story. The stories were manipulated to be either boring or interesting, serving as the “reward” component of the task. Then, in Phase 4, they rated their curiosity again—this time about learning the rest of the story. This reappraisal phase allowed us to examine how story content (prediction error) influenced subsequent curiosity and information-seeking. Finally, in Phase 5, we tested participants’ memory for the photograph pairs and asked them to identify which photograph had been associated with the story they read. Participants’ pupillary responses during PAST were recorded.

**Fig. 2.**
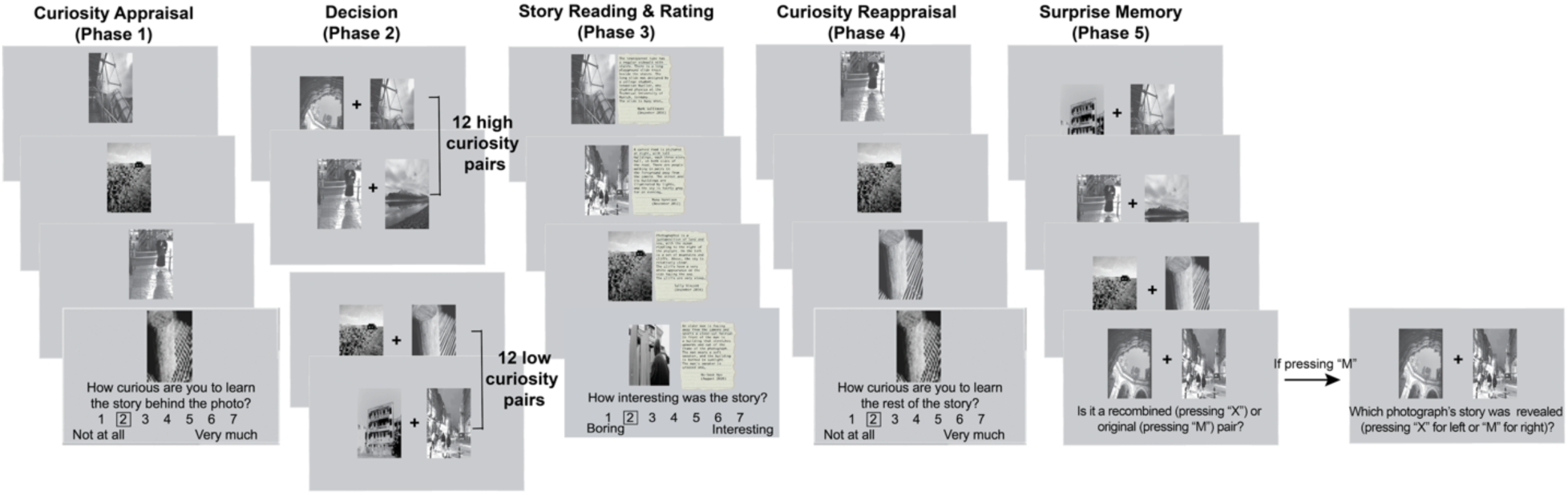
Photographic Art and Storytelling Task. The curiosity-driven information-seeking paradigm leverages curiosity and story outcome to examine the age and individual differences in information-seeking.

Due to poor eye-tracking data quality, 10 young and 13 older adults were excluded from pupillometry analyses but were retained in the behavioral analyses. Moreover, structural MRI data were acquired on a separate day from 43 young adults (using a 7T scanner) and 35 older adults (using a 3T scanner). These scans were used to assess whole-brain and brainstem structures, with a focus on LC-MRI contrast for evaluating LC integrity. To test our hypotheses, we focused on participants’ initial curiosity appraisal (Phase 1), pupil dilation responses during that phase, curiosity reappraisal (Phase 4), and memory performance (Phase 5). We also examined how curiosity-driven information-seeking behavior related to LC structural integrity.

### Age differences in curiosity-induced pupil dilation

We did not observe a significant age difference in initial curiosity ratings during Phase 1 for all photographs (*t*(131) = 1.58, *p* = .12). However, curiosity ratings were differentially associated with task-related pupil dilations across age groups (*p* = .02, 0.82 – 2.04 s; black line in Fig. 3B). In young adults, higher curiosity was linked to greater pupil dilation (*p* = .04, 1.45 – 2.20 s; blue line in Fig. 3B), which is in line with theories that curiosity triggers anticipatory arousal response^26,27^. In contrast, older adults showed a small, inverse relationship (*p* = .049, 0.80 – 1.47 s; orange line in Fig. 3B) with lower curiosity corresponding to slightly larger dilations, perhaps reflecting an aversive response to the less interesting photographs or in response to conflict related to judging photographs negatively. Importantly, this age-related difference was specific to curiosity-induced pupil responses and did not reflect differences in overall pupil size (*p*s > .18; Fig. 3A). These findings suggest that the curiosity-induced pupil dilations may engage different arousal mechanisms across the adult lifespan.

**Fig. 3.**
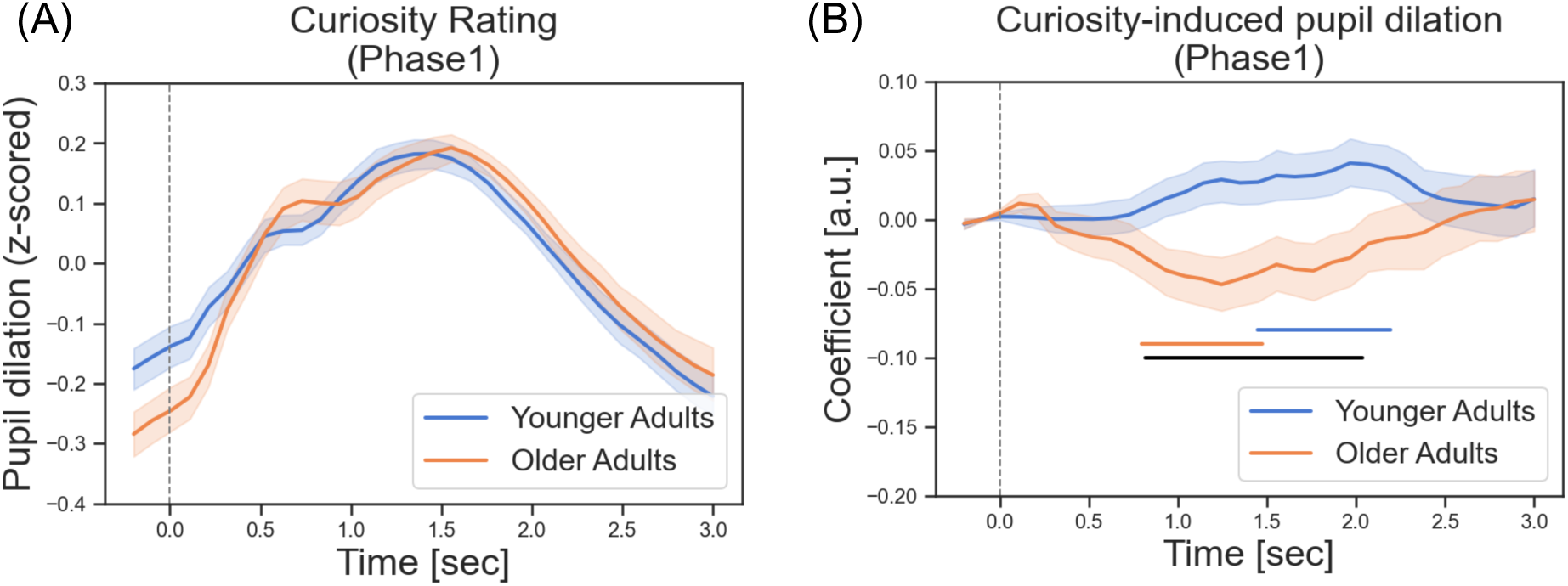
Task-related pupillary responses during curiosity appraisal (Phase 1) in both age groups. (A) No age differences in z-scored pupil dilations during curiosity ratings. (B) The association between curiosity ratings and pupil dilations. Beta regression coefficients accounting for task-related pupil dilations that correlated with trial-by-trial curiosity ratings. Young adults showed a positive association between curiosity ratings and pupil dilations, whereas older adults exhibited a negative association. Blue and orange horizontal lines at the bottom indicate the time points where regression coefficients significantly differentiate from zero in young and older adults, respectively. Black horizontal lines indicate the time points showing significant age differences.

### Curiosity and aging shape the perception of story content and impact further information-seeking

To investigate the extent to which individuals’ information-seeking behavior (i.e., the desire to know the rest of the story during curiosity reappraisal) is driven by initial curiosity (novelty) or influenced by the story content (prediction error), we fit a linear mixed-effects model during curiosity reappraisal. Fixed effects included age group (young vs. older), curiosity context (high vs. low), story outcome (interesting vs. boring), and their interactions. To account for individual differences, we included random intercepts for participants and random slopes for curiosity context and story outcome by participant. Supporting the success of our intended manipulation, interesting story outcomes were associated with higher curiosity ratings upon reappraisal compared to boring ones (main effect of story outcome: B = 2.90, 95% CI [2.63, 3.18], *t* = 20.57, *p* < .001; Fig. 4A). However, initial curiosity judgments from Phase 1 also carried over, such that high-curiosity context photographs maintained higher curiosity ratings upon reappraisal (main effect of curiosity context: B = 1.51, 95% CI [1.24, 1.78], *t* = 10.85, *p* < .001; Fig. 4A). These main effects were qualified by a significant interaction (story outcome × curiosity context: B = -.48, 95% CI [-.76, -.20], *t* = -3.32, *p* = .001; Fig. 4A). Post hoc analyses revealed that the difference between interesting and boring stories was greater in the low-curiosity context (*t* = 2.27, *p* < .001) than in the high-curiosity context (*t* = 1.77, *p* < .001), indicating that curiosity about photographs initially judged as less interesting was more malleable.

**Fig. 4.**
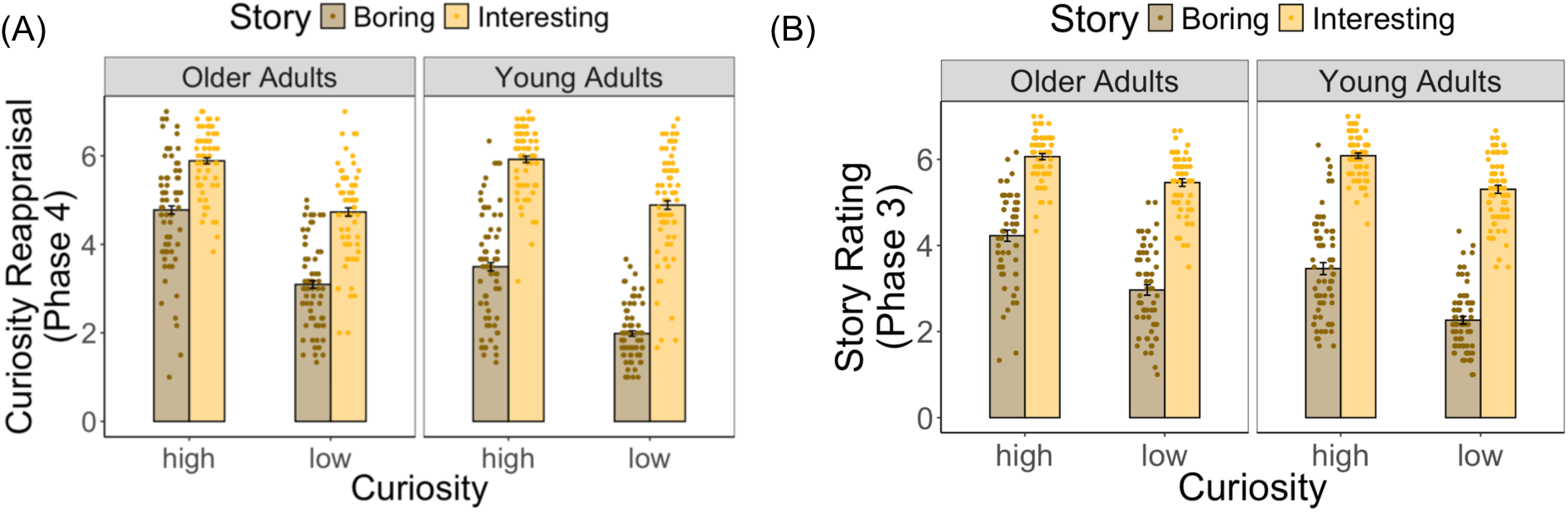
Individual and group ratings for both age groups. (A) Curiosity reappraisal scores, reflecting participants’ desire to know the rest of the story, were shown as a function of curiosity context and story outcome. (B) Story ratings, reflecting how interesting participants found the stories, were also plotted by curiosity context and story outcome.

Critically, we also found age-related differences in information-seeking. Older adults exhibited higher curiosity ratings during reappraisal (main effect of age group: B = 1.11, 95% CI [0.79, 1.43], *t* = 6.88, *p* < .001), but this elevation appeared to reflect reduced updating based on story outcomes (story outcome × age group: B = -1.27, 95% CI [-1.66, -.87], *t* = -6.28, *p* < .001; Fig. 4A). Specifically, the difference between interesting and boring stories was smaller in older adults (*t* = 1.37, *p* < .001) than in young adults (*t* = 2.66, *p* < .001), suggesting that older adults rely more on their initial levels of curiosity, while young adults’ information-seeking is more likely to be updated and influenced by story outcomes. No other significant interactions were observed (*ps* > .39).

One concern was that age-related memory decline, as shown in the memory performance results below, may have influenced curiosity reappraisal, with older adults relying more on initial curiosity than on the actual story content due to forgetting details of the stories. To test this, we conducted the same analysis on participants’ ratings of how interesting the stories were during the reading phase. The results showed consistent main effects of age group (*B* = .70, 95% CI [.40, 1], *t* = 4.57, *p* < .001), curiosity context (*B* = 1.20, 95% CI [.99, 1.41], *t* = 11.06, *p* < .001), and story outcome (*B* = 3.04, 95% CI [2.78, 3.31], *t* = 22.57, *p* < .001). We also observed significant interactions between curiosity context and story outcome (*B* = -.42, 95% CI [-.67, -.16], *t* = -3.23, *p* = .001; Fig. 4B) and between age group and story outcome (*B* = -.55, 95% CI [-.92, -.17], *t* = -2.84, *p* = .005; Fig. 4B). Post hoc analyses supported the same pattern, showing that the difference between interesting and boring stories was greater in the low-than in the high-curiosity context (*t* = 2.23, *p* < .001). The difference in story ratings between interesting and boring stories was larger in young adults (*t* = 2.83, *p* < .001) than in older adults (*t* = 2.17, *p* < .001). Together, these findings suggest that aging and individuals’ initial curiosity influence the perception of story contents and affect subsequent information-seeking behavior.

### Locus coeruleus integrity predicts bias toward novelty-driven information seeking

We found a lower LC-MRI contrast ratio in older adults compared to young adults (*t*(76) = -5.04, *p* < .001; Fig. 5A), suggesting age-related decline in LC structural integrity^15,28^. To better understand the influence of story content on individuals’ information-seeking behavior, we calculated a “story bias ratio,” which captures the change in curiosity ratings from the initial appraisal (Phase 1) to the reappraisal after reading the story (Phase 4), normalized by the sum of these two ratings. A higher story bias ratio indicates that the story content had a greater effect on curiosity reappraisal, reflecting prediction error-based curiosity. In contrast, a lower bias ratio suggests that curiosity reappraisal remained similar to the initial ratings, indicating novelty-driven curiosity. We observed substantial rating changes only for photographs whose associated stories were read. In contrast, curiosity ratings for photographs without story exposure remained largely unchanged (see Fig. S2 in Supplementary Materials), suggesting that the story content influenced participants’ curiosity. Building on this behavioral effect, we fit a linear regression model with the story bias ratio as the dependent variable. The model included LC-MRI contrast ratios, curiosity context, story outcome, age group, and their interactions as predictors, while controlling for chronological age, sex, and years of education. This analysis tested whether variability in LC integrity was associated with individual differences in the balance between novelty- and prediction error-driven information seeking. The results revealed that participants with a higher LC-MRI contrast ratio, reflecting greater LC structural integrity, exhibited a lower story bias ratio (*B* = -0.30, 95% CI [-0.59, -0.01], *t* = -2.05, *p* = .04; Fig. 5B). This effect did not interact with age group (*B* = 0.12, 95% CI [-0.49, 0.73], *t* = -0.40, *p* = .69), indicating that the relationship between LC integrity and reduced reliance on story content was consistent across young and older adults. Given the high correlation between age group and MRI field strength (which differed across age cohorts), we reran the model, excluding age group, and instead included field strength as a covariate to account for differences in imaging modalities. The pattern of results remained unchanged: individuals with greater LC structural integrity consistently demonstrated a lower story bias (*B* = -0.27, 95% CI [-0.53, -0.02], *t* = -2.14, *p* = .04). These findings suggest that greater LC integrity may bias individuals toward novelty-driven, rather than prediction error–based, information-seeking.

**Fig. 5.**
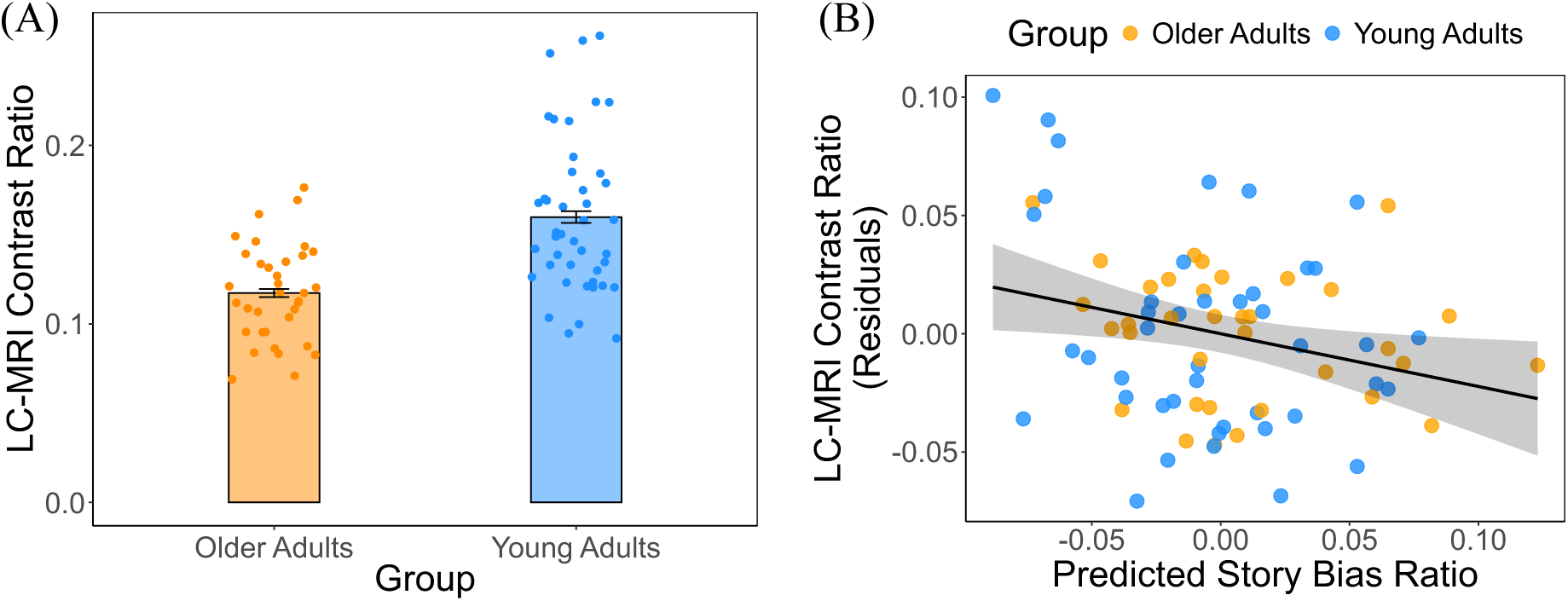
Locus coeruleus (LC) structural integrity and its association with information-seeking behavior. (A) Age-related decline in the LC structural integrity. (B) A negative association between the LC structural integrity and story bias ratio, suggesting a role of the LC in regulating curiosity-driven information-seeking.

### Curiosity enhances memory performance

Curiosity’s influence on memory performance was evaluated using a linear mixed-effects model, with age group and curiosity context entered as fixed effects and random intercepts specified for participants. A significant age-related difference emerged in recognition memory, which was tested during Phase 5 (*B* = -0.14, 95% CI [-0.17, -0.10], *t* = -6.76, *p* < .001; Fig. 6A), where participants judged whether paired photographs had been previously presented during the decision phase. There was no significant main effect of curiosity context (*B* = 0.01, 95% CI [-0.01, 0.04], *t* = 0.91, *p* = .37), nor a significant interaction between age group and curiosity context (*B* = -0.01, 95% CI [-0.05, 0.03], *t* = -0.58, *p* = .57). However, a curiosity-related enhancement was evident in the subsequent memory test, specifically in participants’ ability to identify which photograph whose story had been revealed after endorsing that the pair had been shown previously (*B* = 0.03, 95% CI [0.003, 0.06], *t* = 2.17, *p* = .03; Fig. 6B). While an overall age-related difference in this subsequent memory performance was observed (B = -0.11, 95% CI [-0.14, -0.07], t = -5.54, *p* < .001; Fig. 4B), the absence of a significant interaction between age group and curiosity context (B = 0.02, 95% CI [-0.01, 0.06], t = 1.16, *p* = .25; Fig. 6B) revealing consistent beneficial effects of curiosity on memory in both young and older adults.

**Fig. 6.**
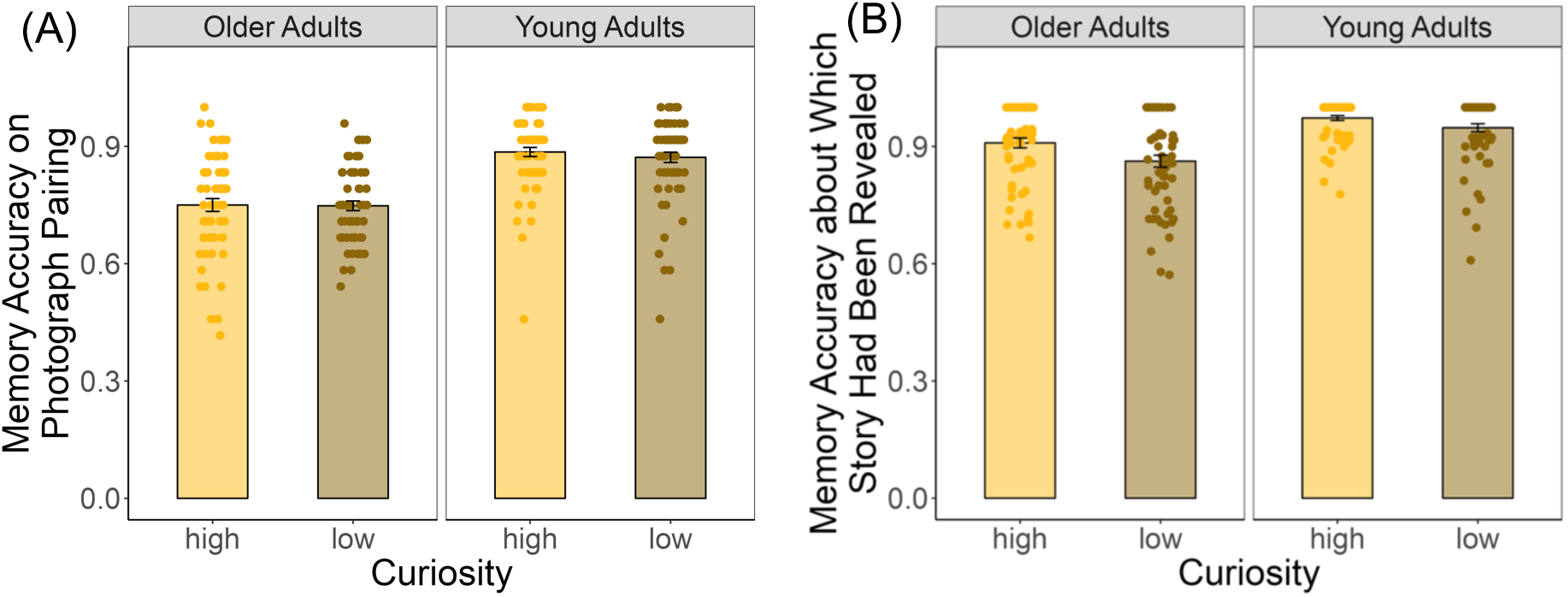
Individual and group memory performance for both age groups. (A) Memory performance about whether paired photographs had been previously presented during the decision phase. (B) Memory performance about which photograph’s story had been revealed during story reading phase.

To examine whether individual differences in LC integrity contributed to this curiosity-related memory boost, we conducted an exploratory analysis including LC-MRI contrast ratios. Results showed no main effect of LC integrity (*B* = -0.12, 95% CI [-0.39, 0.15], *t* = -0.90, *p* = .37), nor any significant interactions with age group or curiosity context (*p*s > .44; see Supplementary Materials for details). Taken together, these findings indicate that curiosity can support memory for actively encoded information across the adult lifespan and may serve as a motivational factor to maintain cognition in aging.

## Discussion

This study employed a novel paradigm that engaged both novelty-driven and prediction error-based curiosity to examine how curiosity and aging influence information-seeking behavior, as well as their relationship with LC integrity. Our findings revealed that young adults adapt their information-seeking based on story content, which acts as a rewarding context involving prediction error, whereas older adults rely more heavily on novelty-driven (initial) curiosity. We also observed age-related differences in curiosity arousal and engagement, as reflected in pupillary responses. Moreover, individual differences in LC integrity were associated with a shift away from the story-based (prediction error-based) effect on guiding information-seeking behavior. Finally, we found that curiosity enhances memory performance, particularly for information that is active (which photograph’s story was revealed), across the adult lifespan. These findings offer new insights into the role of the LC-norepinephrine system in curiosity-driven information-seeking in aging.

To create the curiosity contexts (high vs. low), we encouraged participants to use the full range of the rating scale. This approach may have contributed to the lack of observed age differences in the initial curiosity ratings. However, in the subsequent information-seeking behavior (i.e., knowing the rest of the stories), older adults demonstrated higher curiosity compared to young adults. This finding aligns with previous studies using a trivia question paradigm, which similarly reported increased curiosity-driven behavior in older adults^29–31^. This heightened curiosity may reflect age-related motivational shifts toward acquiring meaningful or personally relevant knowledge^30,32^. Given that curiosity has been linked to heightened arousal and attentional engagement, we next examined pupil dilation as a psychophysiological index of these processes across age groups. In young adults, pupil dilations positively correlated with curiosity ratings, consistent with previous research suggesting that curiosity enhances anticipatory arousal in response to upcoming information^26,27^. In contrast, the relationship between curiosity and pupil dilation was more complex in older adults. Specifically, we observed a small inverse relationship, where lower curiosity ratings were associated with larger pupil dilations. This may indicate that older adults experienced an aversive or disengaged response to less interesting stimuli^33,34^, possibly reflecting age-related changes in attentional or affective processing. Another possibility is that older adults’ positivity bias^35,36^ led to a general readiness to engage with the stories, such that assigning low curiosity scores conflicted with this tendency and elicited heightened arousal. Overall, these findings suggest that while pupil dilation remains a valuable marker of curiosity-related arousal, its interpretation may differ with age due to underlying cognitive and emotional shifts.

Contrary to the trivia paradigm commonly used in curiosity research^30,31,37–39^, which often allows participants to draw on prior knowledge, our PAST paradigm introduces a novelty component and context in which participants have no prior experience, engaging the LC-norepinephrine system. This design provides a unique opportunity to examine how aging influences information-seeking behavior, particularly whether it shifts toward novelty-driven versus prediction error-based curiosity. Indeed, we found that story content influenced participants’ subsequent information-seeking behavior, but only for the photographs whose associated stories were revealed; no such effect was observed for photographs whose stories were not read (see Fig. S2). Importantly, young adults’ information-seeking behavior was more strongly shaped by story content, suggesting a reward-related prediction error mechanism linked to the dopaminergic system. In contrast, older adults appeared to rely more on their initial feelings of curiosity, indicating a greater sensitivity to novelty and potential involvement of the norepinephrine system. These findings suggest that young and older adults engage in qualitatively different strategies for information-seeking, potentially reflecting age-related shifts in underlying neuromodulatory systems.

Beyond age-group differences in information-seeking approaches, we also found evidence for individual variability: participants’ initial curiosity interacted with story content to influence subsequent information-seeking behavior. Specifically, the impact of story content was more pronounced for photographs that initially elicited low curiosity compared to those that evoked high curiosity. This suggests that early curiosity judgments can anchor subsequent engagement. The findings further support the idea that both external rewards and initial curiosity shape information-seeking behavior, consistent with the dual neuromodulatory system framework^1,5^ (Fig. 1). Moreover, the extent to which story content influenced information-seeking was negatively associated with LC integrity, supporting the idea that the LC-norepinephrine system modulates curiosity-driven exploration^6–8^. Altogether, the results point to both age-related differences and individual variability in how people shift along the continuum between prediction error–based and novelty-driven curiosity, and how this variability may relate to the LC–norepinephrine system.

In addition to guiding information-seeking behavior, curiosity also plays a critical role in enhancing memory. This motivational state not only drives exploration but also facilitates deeper encoding of information. Our findings showed that curiosity enhanced memory performance, but only when participants actively engaged with and encoded the information (which photograph’s story was revealed), indicating that curiosity’s mnemonic benefits depend on intentional, goal-directed processing^38,39^. Notably, this curiosity-related memory enhancement was observed in both young and older adults, suggesting that the beneficial effects of curiosity on learning are preserved across the adult lifespan. This aligns with prior work demonstrating that states of high curiosity engage neuromodulatory systems, such as the dopaminergic and norepinephrinergic systems, which in turn support hippocampus-dependent memory formation^1,5^. Although we found no direct relationship between curiosity-driven memory enhancements and LC structural integrity (see details in Supplementary Materials), it is possible that curiosity facilitates memory formation via mesolimbic functional connectivity associated with these neuromodulatory systems^38,40,41^. Future studies could administer functional neuroimaging to explore how curiosity influences mesolimbic pathways involved in memory encoding and consolidation. Together, these results emphasize that while aging may alter how curiosity drives information-seeking, the capacity for curiosity to enhance learning remains relatively intact in older age, which indicates that curiosity may serve as a motivational factor to support cognitive resilience in aging^42^.

There are several limitations to the current study. First, our sample size was relatively small, and the observed effects were based on cross-sectional comparisons between young and older adults. While this approach provides initial insights into age-related differences, it does not capture developmental trajectories across the lifespan. Of note, recent research has identified a quadratic pattern in state curiosity across age, with older adults showing higher curiosity than middle-aged adults^30^. This nonlinear trend aligns with evidence that neuromodulatory systems, such as the LC-norepinephrine system, also follow an inverted-U trajectory with age^16–18^. These patterns suggest that both curiosity and its underlying neural mechanisms may change in complex, interactive ways over the lifespan. Future studies employing longitudinal designs and broader age ranges are needed to better understand how aging and neuromodulatory function jointly shape curiosity-driven learning and exploration. Second, although we speculated that young adults’ information-seeking behavior is more strongly driven by prediction errors, potentially linked to the dopaminergic system, we did not collect psychophysiological or neural measures to directly support this claim. Previous research has shown that pupil dilation is sensitive to prediction errors^42,43^, suggesting that pupil size could serve as an indirect index of such processes. One possible approach in our study would have been to examine pupil dilation during the story reading phase. However, this method presents significant challenges: natural reading involves frequent eye movements and varying gaze angles, both of which can distort pupil size measurements. In addition, we found substantial individual and age differences in reading time (see Fig. S2 in the Supplementary Materials), introducing further variabilities that complicate the interpretation of pupil data. These factors limit our ability to assess prediction error-related arousal through pupillometry in the current paradigm. To more directly investigate the neural mechanisms underlying prediction error-based information seeking, future studies could assess the structural integrity of dopaminergic midbrain regions, such as the substantia nigra (SN) or the ventral tegmental area (VTA)^44,45^. These regions are known to support reward processing and learning and could provide more direct evidence linking age-related differences in dopaminergic function and curiosity-driven information-seeking.

To conclude, our study revealed age-related differences in information-seeking behavior. Young adults’ information-seeking was more influenced by prediction-error-related curiosity, whereas older adults relied more heavily on novelty-driven curiosity. Individuals with higher LC integrity exhibited greater novelty-driven curiosity. These findings highlight the impact of aging and the role of the LC in modulating curiosity-driven information-seeking. Importantly, curiosity enhanced memory when the information was actively encoded—a benefit that was preserved in both young and older adults. The present study further showed that curiosity continues to play a vital role in motivating learning and exploration across the adult lifespan, with distinct expressions at different ages. While young adults flexibly updated their curiosity in response to outcomes, older adults showed a tendency to rely more on their initial judgments. This pattern may reflect a shift in how curiosity guides exploration: older adults may prioritize what initially captures their interest, while young adults remain more sensitive to prediction errors. Both pathways, however, can support engagement and learning, with implications for lifelong exploration, cognitive health, and the maintenance of curiosity-driven motivation into older adulthood.

## Methods

### Participants

A total of 133 participants (68 young and 65 older adults) were recruited in this study. Table 1 depicts participants’ demographic information. All participants reported normal or corrected-to-normal vision and no history of psychiatric or neurological disorders. To control for factors known to influence pupil dilations, participants were instructed to abstain from alcohol for 24 hours prior to the pupillometry session and to refrain from consuming caffeinated beverages (e.g., coffee or tea) on the day of testing^46^. All experimental sessions were conducted between 9:00 AM and 5:00 PM to minimize the potential influence of circadian fluctuations on pupil dilation^47^. Data from 10 young and 13 older adults whose eye-tracking data quality was poor (> 50% of the data needed to be removed from the analyses of pupil size) were included in the behavioral analyses but excluded from the pupillometry analyses. A subset of participants, consisting of 43 young and 35 older adults, underwent structural MRI using either a 7T or 3T scanner, respectively, on a separate day to acquire whole-brain and brainstem images for assessing the LC-MRI contrast. Written informed consent in accordance with the Declaration of Helsinki (2008) was obtained from all participants. This study was approved by the Brandeis University Human Research Protection Program and the Mass General Brigham Human Research Committee.

**Table 1.**
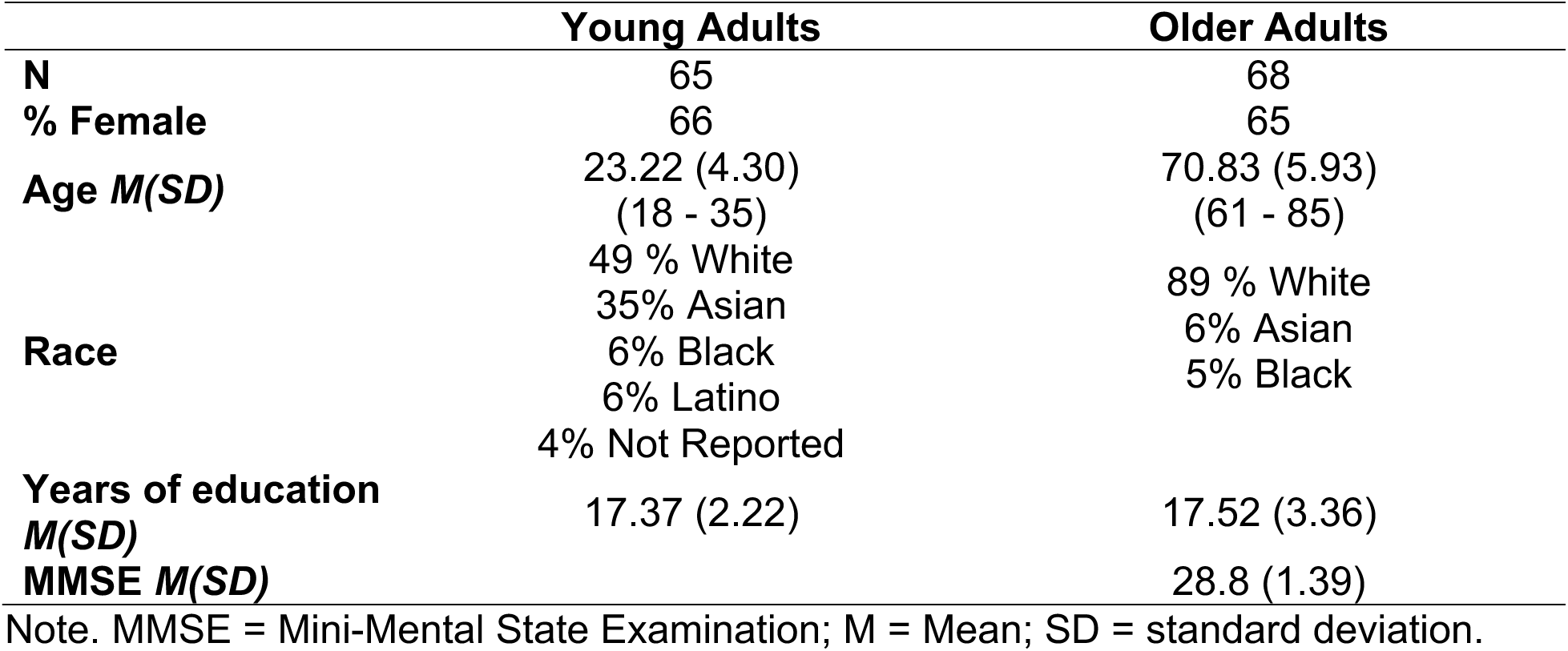
Participant Demographics.

### Photographic art and storytelling task

The PAST was inspired and adapted from Biderman and Shohamy (2021)^48^, which used monetary rewards (extrinsic value) rather than curiosity (intrinsic motivation) manipulations to probe interactions among reward, decision-making, and memory in young adults. Fig. 1 depicts the PAST paradigm. The task involves five phases: 1) curiosity appraisal, 2) decision, 3) story reading (context-based reward), 4) curiosity reappraisal, and 5) surprise memory.

#### Curiosity appraisal (Phase 1)

Participants viewed 88 black-and-white photographs, each accompanied by information that each photograph had a story behind it. Each photograph was presented for 3 seconds, during which no responses were permitted. This fixed 3-second viewing period was designed to capture pupillary responses associated with curiosity. After the 3-second interval, a 7-point rating scale appeared below the image, prompting participants to indicate their level of curiosity, specifically, how strongly they desired to learn the story behind the photograph. To help generate meaningful curiosity levels for subsequent phases, participants were encouraged to use the full range of the rating scale.

#### Decision (Phase 2)

Individual participants’ ratings in Phase 1 were sorted and formed into 12 high- and 12 low-curiosity pairs. Participants were asked to choose one photograph from each pair to reveal the story (each pair was repeated three times). This phase established the curiosity condition (high vs. low) and facilitated memory encoding for later recall in the memory test.

#### Story reading (Phase 3)

The first part of each selected story was revealed. The stories varied in their level of information richness (interesting vs. boring), which creates value as a reward. Interesting stories offer new details about a person’s experience or remarkable history, while boring stories only provide basic descriptions of the photographs. Each story consisted of three sentences (interesting: M ± SD = 50.31 ± 3.99 words; boring: M ± SD = 52.20 ± 3.86 words). Participants were instructed to rate how interesting the story was after reading.

#### Curiosity reappraisal (Phase 4)

Participants rated their curiosity levels again, reflecting their desire to know the rest of the story (subsequent information-seeking). This phase allows us to examine whether further information-seeking (e.g., knowing the rest of the story) is driven by novelty-driven curiosity, where individuals maintain curiosity levels regardless of whether the story is perceived as interesting or boring, or by prediction error-based curiosity, where curiosity shifts based on the perceived interest or appeal of the story.

#### Surprise memory (Phase 5)

Participants were initially asked to determine whether each presented pair of photographs had previously been shown together during the decision phase (Phase 2). For pairs identified as the same, participants were subsequently required to indicate which photograph’s story had been revealed.

The task was programmed in MATLAB R2022b (the MathWorks, Inc, 2015) with the Psychtoolbox software^49^. Each trial in each phase started with a fixation cross at the center of the screen with a mean duration of 1,500 ms, followed by a stimulus. Stimuli were displayed on a 24-inch monitor (1920 × 1080 pixels) and viewed from a distance of 60 cm. The photographic stimuli in Phases 1 and 4 were presented at a size of 7 × 11 cm at the center of the screen, whereas the stimuli in Phases 2, 3, and 5 were presented at a size of 15 × 11 cm. The luminance of the photographs shown on the screen was adjusted (M ± SD = 72.04 ± 0.16 cd/m^2^) using a Konica Minolta LS-150 luminance meter to control for potential luminance-related effects on pupil size. The head position of the participants was stabilized using a chin and forehead rest.

### Acquisition and preprocessing analysis of pupillometry data

Participants’ pupil size was continuously recorded during the task using an EyeLink 1000 plus eye-tracker (SR Research, Canada) with a sampling rate of 1000 Hz. A 9-dot eye-tracking calibration was conducted at the beginning of each task phase. We focused on the curiosity-induced pupil dilations during curiosity appraisal (Phase 1). To this end, pupillometry data were segmented from 200 ms before and 3,000 ms after the photograph was shown. A customized Python script detected eye-blinks in the segmented data. The periods of missing data due to blink artifacts were segmented in a time window of 100 ms before and 100 ms after the blink and replaced by a cubic interpolation. As such, the interpolation was only applied to periods where data loss durations were shorter than 750 ms^43,50^. Trials with excessively noisy or missing data in which the blink artifacts sustained over 750 ms were excluded within subjects (removed trials: M ± SD = 0.09 ± 0.11 in young and 0.1 ± 0.1 in older adults). Pupillometry data were then baseline corrected with regard to the first 200 ms of each segmented time window, and standardized *z*-scores were calculated within participants to allow comparing pupil dilations associated with curiosity ratings independent of individual differences in mean and variance of pupil size^43,51^.

### Acquisition of structural MRI data, LC-MRI contrast evaluation, and gray matter volume analysis

Older adults’ brain images were obtained from a longitudinal Brandeis Aging Brain Study. Structural MRI scans were acquired on a 3T Prisma whole-body scanner (Siemens, Erlangen, Germany) with a standard 64-channel head coil used for signal reception. A high-resolution T1-weighted anatomical image was acquired using a multi-echo magnetization-prepared rapid acquisition gradient echo (MEMPRAGE) sequence (voxel size = 1 × 1 × 1 mm, 256 slices, TR = 2,530 ms, TEs = [1.69, 3.55, 5.41, 7.27] ms, flip angle = 7°, field-of-view [FOV] = 256 mm, acquisition matrix = 176 × 256). To visualize the LC, a 2D T1-weighted turbo-spin-echo (TSE) sequence with additional magnetization transfer (MT) contrast was applied (voxel size = 0.4 × 0.4 × 3 mm, six slices, TR = 743 ms, TE = 15 ms, flip angle = 180°, four online averages, and acquisition time = 3 min and 22 s)^16^. Thirteen participants completed a single LC scan, while twenty-two underwent two LC scans. No significant differences were observed between participants who completed one scan and those who completed two (B = -0.01, 95% CI [-0.02, 0.01], *t* = - 1.10, *p* = .28; Fig. S3), with further details provided in the Supplementary Materials.

Young adult participants’ structural MRI scans were acquired on a 7T Terra whole-body scanner (Siemens, Erlangen, Germany) with a 32-channel head coil. A high-resolution whole-brain image was acquired using a magnetization prepared 2 rapid acquisition gradient echoes (MP2RAGE) sequence (voxel size = 0.8 × 0.8 × 0.8 mm^3^, 240 slices, TR = 4,000 ms, TE = 1.92 ms, flip angles = [5°, 6°], FOV = 240 mm, acquisition matrix = 192 × 210). In addition, two 3D magnetization transfer-weighted turbo flash (MT-TFL) scan, developed by a previous study^52^, was performed to scan the LC (voxel size = 0.4 × 0.4 × 0.5 mm^3^, 60 slices, TR = 400 ms, TE = 2.55, flip angle = 8°, acquisition time = 8 min and 13 s). For the MT-TFL sequence, the FOV was oriented approximately perpendicular to the pons and spanned the region from the superior colliculus to the mid-pons border.

Signal intensities in the LC region were extracted from each participant’s TSE-MT or MT-TFL image using a binarized LC mask previously published^25^. Brainstem image registration proceeded in two main steps^43^. First, each participant’s brainstem images were aligned to their native whole-brain images (Step 1). Data from participants who underwent two LC-MRI scans were then averaged for subsequent analysis. Second, the whole-brain images were co-registered to MNI 0.5 mm linear space, in which the LC mask was defined, to obtain the transformation matrices (Step 2). Both registration steps used the “antsRegistration” function from Advanced Normalization Tools^53^ (ANTs, version 2.1). Step 1 involved linear registration (rigid followed by affine), while Step 2 incorporated additional non-linear registration (SyN) to improve alignment with the MNI template. The transformation matrices obtained from Step 2 were then applied, in reverse, to warp the LC mask from MNI space back into each participant’s native brainstem image space using nearest-neighbor interpolation. A pontine reference mask consisting of two bilateral 4 × 4 mm regions was created for each participant. The dorsal boundary of this reference region was defined by shifting 8 mm ventrally from the dorsal edge of the LC region into the pons^25^. These masks were then used to extract signal intensities. To allow normalization and between-participant comparisons, the LC-MRI contrast ratio was calculated on each slice along the rostrocaudal axis of the LC using the following formula:

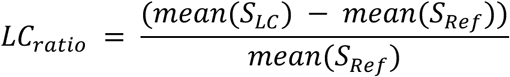

where *mean(S_LC_)* denotes the mean signal intensity of the left or right LC, and *mean(S_Ref_)* indicates the mean signal intensity of the pontine reference region in the corresponding slice. In other words, the LC signal intensity was standardized using nearby white matter tissue. To obtain more stable intensity estimates for subsequent correlation analyses, LC contrast ratios were averaged across hemispheres and slices along with the LC dimension within each participant.

### Statistical analysis

Participants’ curiosity ratings (Phases 1 and 4), story ratings (Phase 3), and memory accuracies (Phase 5) were analyzed using multi-level (linear mixed) models in R (version 4.3.1). To assess the extent to which participants’ subsequent information-seeking behavior (knowing the rest of the story in Phase 4) was modulated by the content of the story, a story bias ratio was computed. This ratio was defined as the absolute difference between participants’ initial curiosity appraisal and their reappraisal for the same photograph after reading the associated story, divided by the sum of the curiosity ratings before and after reading the story: |Phase 4 – Phase 1| / (Phase 4 + Phase 1). This measure captures the proportional change in curiosity, accounting for individual differences in overall curiosity levels. Multi-level modeling was performed on the behavioral measures described above using the “lme4” package (version 1.1-34). Post hoc comparisons were conducted using the “emmeans” package (version 1.8.8), with Bonferroni correction applied at a significance level of p < 0.05.

To examine curiosity-induced pupil dilation, trial-by-trial curiosity ratings from Phase 1 were correlated with pupil dilation values at each time point within a 3-second window following the image presentation, using the linear regression model in the “sklearn” package (version 1.6.1) in Python (version 3.10.16). To examine age differences in curiosity-related pupil responses, non-parametric cluster-based permutation t-tests were conducted using the MNE package (version 1.9.0). The cluster size was determined by the number of continuous time series for which the *t*-test resulted in *p* < 0.05, and then the observed cluster size was compared against a random cluster across subjects within or between age groups (5,000 repetitions) to quantify the significance of this cluster in the time series.

Furthermore, linear regression models were used to test whether individuals with greater LC structural integrity exhibited lower story bias ratios, indicating that their information-seeking behavior was more strongly driven by novelty-related curiosity. We also explored the associations between the LC structural integrity and task-related pupil dilations and between the LC structural integrity and memory performance. Linear regression modeling using the “stats” package (version 4.3.1) in R was applied to examine the relationships between brain metrics and behavioral measures. Chronological age, sex, and years of education were included as covariates.

## Acknowledgments

We thank all participants for taking part in the study. We are especially grateful to Sinead Chen, Grace, and a close friend for providing the photographic materials used. We appreciate Jennifer L. Crawford for her valuable input and comments in support of this work. We also acknowledge Megan Applegate-Kenton for her assistance with project administration. This work was supported by R00AG058748 (ASB); R01AG074330 (ASB); R21AG086729 (ASB); Scientific Research Network on Decision Neuroscience & Aging – Pilot grant (HYC).

## Author contributions

A.S. Berry and H.-Y. Chen conceived the research idea and designed the study. K.E. O’Malley and H.-Y. Chen wrote the associated stories used in the task. H.-Y. Chen, E.L. Carlson, M.S. Costello, J.L. Matulonis, and A.A. Adornato contributed to data collection. H.I.L. Jacobs and J.M. Hooker provided guidance on neuroimaging testing. H.-Y. Chen made contributions to the analysis and interpretation of data. A.S. Berry and H.-Y. Chen drafted the manuscript. A.S. Berry and H.-Y. Chen performed the writing review and editing, and all authors read and approved the final version of the manuscript. A.S. Berry and H.-Y. Chen supervised the study.

## Data availability statement

All de-identified behavioral data and analytic code are publicly available on the Open Science Framework (https://osf.io/qw4j2/?view_only=7b1540d4a53e46ffa614b8087699c848).

