## Supplementary material for "Age-Related Changes in Curiosity: The Influence of Locus Coeruleus on Information-Seeking Behavior": Supplmentary materials

### Supplementary Materials

#### Individual and age differences in response time during the story reading phase

Furthermore, we also observed substantial individual and age-related differences in response times during the story reading phase, which complicate the analysis of pupil dilations in this part of the task. Table S1 and Fig. S1 display the distributions of response times across task conditions and age groups. Specifically, older adults took significantly longer to read the stories than younger adults (B = 3.30, 95% CI [1.59, 5.02], $t = 4.06$, $p < .001$). Participants also spent more time reading stories associated with high-curiosity photographs compared to low-curiosity ones (B = .77, 95% CI [.06, 1.48], $t = 2.50$, $p = .01$), and more time reading stories they rated as interesting rather than boring (B = 1.44, 95% CI [.73, 2.15], $t = 3.72$, $p < .001$). Although the story reading phase involved prediction error processing (e.g., high curiosity photograph associated with boring story and vice versa), these variations in reading duration pose a challenge for investigating pupil dilations during the story reading phase, as the considerable variability in the timing and length of cognitive engagement across participants makes it difficult to temporally align pupillary responses in a consistent and interpretable manner.

**Table S1.** Means and standard deviations of response time during the story reading phase in each task condition and each age group.

| | Young Adults | | Older Adults | |
|---|---|---|---|---|
| | Interesting Story | Boring Story | Interesting Story | Boring Story |
| High Curiosity | 13.57 ± 5.51 | 12.42 ± 4.85 | 16.58 ± 5.54 | 15.49 ± 4.56 |
| Low Curiosity | 13.09 ± 4.83 | 11.64 ± 4.43 | 16.59 ± 5.47 | 14.95 ± 5.19 |

Note. Response times in seconds.

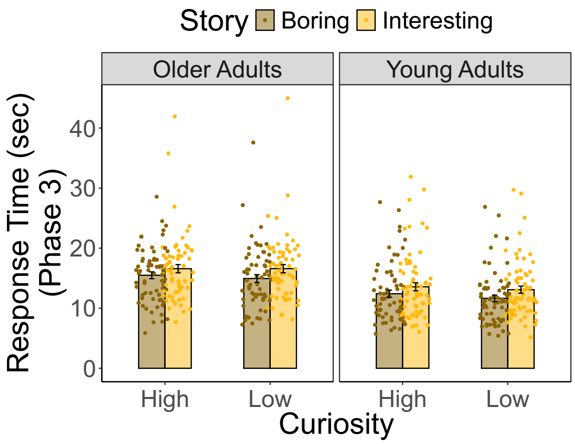


**Fig. S1. Response time about the story rating during reading in both age groups.**

### Curiosity ratings remain stable for photographs without associated story exposure

To determine whether story content influenced participants' ratings only for photographs whose associated stories were revealed, we also calculated the rating change ratio (the change of curiosity ratings between Phase 1 and Phase 4) for photographs with unrevealed stories. This ratio was defined as the absolute difference between participants' initial curiosity appraisal (Phase 1) and their reappraisal of the same photograph (Phase 4), divided by the sum of the two curiosity ratings. We then conducted a linear regression analysis with the rating change ratio as the dependent variable, and story status (revealed vs. unrevealed), age group, and their interaction as predictors. The results showed a significant main effect of story status ($B$ = .17, 95% CI [.16, .19], $t$ = 22.99, $p$ < .001; Fig. S2), as well as a significant interaction between age group and story status ($B$ = -.03, 95% CI [-.04, -.005], $t$ = -2.42, $p$ = .02; Fig. S2). There was no significant main effect of age group ($B$ = .001, 95% CI [-.01, .02], $t$ = .18, $p$ = .85). These results indicate that only the photographs whose stories were read showed larger rating changes compared to those whose stories were not revealed. Post hoc analysis further revealed that, for photographs with revealed stories, young adults exhibited a greater change ratio than older adults ($t$ = 3.24, $p$ = .001).

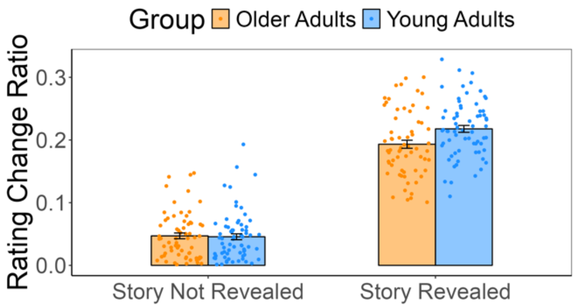


**Fig. S2. Ratios of rating changes between curiosity appraisal and reappraisal phases in both age groups.** The curiosity ratings changed more in the photographs whose stories were read, whereas the ratings remained consistent in those whose stories were not revealed.

### LC-MRI contrast ratios do not differ significantly between participants with one or two MT-TSE scans

Because some older adults underwent one MT-TSE scan while others had two, we used a linear regression model to test for differences, including the number of scans as a fixed effect and controlling for chronological age, sex, and years of education. The results showed no significant effect of scan number (B = -.01, 95% CI [-.02, .01], $t$ = -1.10, $p$ = .28), indicating that the average LC-MRI contrast within subjects is reliable for further analyses.

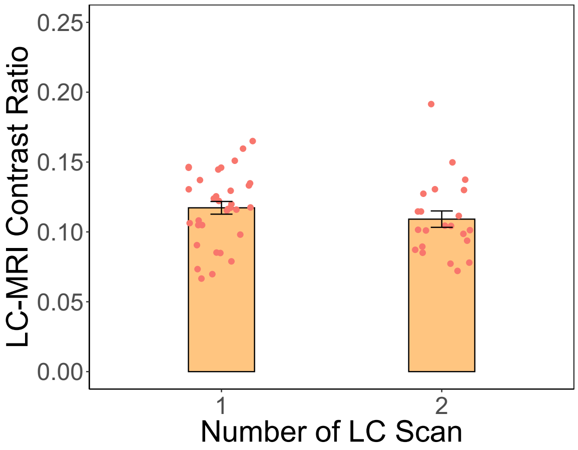


**Fig. S3. LC-MRI contrast ratio in participants who had one or two MT-TSE scans.**

**No association between LC integrity and curiosity-driven effects on memory performance**

We investigated whether individual differences in LC integrity predict curiosity-driven effects on memory performance. A linear regression model was conducted with memory accuracy for photographs whose accompanying stories had been revealed as the dependent variable. LC-MRI contrast ratio, age group, and curiosity context were included as fixed effects, with random intercepts specified for participants. Chronological age, sex, and years of education were included as covariates. The results revealed no significant main effect of LC-MRI contrast ratio ($B$ = -0.12, 95% CI [-0.39, 0.15], $t$ = -0.90, $p$ = .37), nor any significant interactions with age group or curiosity context ($p$s > .44).
